# Gut Microbiome Signatures Reflect Within-Host Subtype Diversity of *Blastocystis* in Non-Human Primates

**DOI:** 10.1101/2025.02.25.640083

**Authors:** Pingping Ma, Wenjie Mu, Yugui Wang, Yihui Liu, Yang Zou, Hong Pan, Long Zhang, Lixian Chen, Yongpeng Yang, Weifei Luo, Shuai Wang

## Abstract

*Blastocystis* is a prevalent gut eukaryote intricately associated with the gut microbiota. This genetically diverse protozoan exhibits significant intra-host subtype heterogeneity, yet the implications of this diversity for the host gut microbiome remain poorly understood. Here, we investigated the interactions between *Blastocystis* and gut microbiota in non-human primates at the level of subtypes, using a comprehensive investigation of gut microbiota for *Blastocystis* carriers of captive *Macaca fascicularis* (discovery cohort, n = 100) and *Macaca mulatta* (validation cohort, n = 26). We identified highly prevalent intra-host co-occurrence patterns of *Blastocystis SSU* rRNA-based subtypes, primarily dominated by Subtype 1(ST1) or ST3. These patterns were associated with compositional and structural variations in the gut microbiome but were not significantly influenced by host covariates such as sex, age, or BMI. Specifically, Ruminococcaceae-enterotype was enriched in the patterns dominated by ST1, whereas *Limosilactobacillus*-enterotype was predominantly identified in the patterns dominated by ST3. We discovered that the absolute abundance of *Blastocystis* was a critical determinant in elucidating this association across concurrent patterns. *In vivo* experiments in a new cohort (n = 11) demonstrated that lactic acid bacteria, enriched in the *Limosilactobacillus*-enterotype, were sufficient to reduce *Blastocystis* load. We validated the strong association between gut microbiome composition and *Blastocystis* load in *M. mulatta*, confirming that specific microbial features could quantitatively predict *Blastocystis* status in both species. Our findings establish a direct link between gut microbial variations, intra-host genetic heterogeneity, and absolute abundance in *Blastocystis* for non-human primates.

## Introductions

*Blastocystis* is a widely prevalent eukaryotic microorganism inhabiting the gut of both humans and animals^1^. It belongs to the heterokonts clade, which encompasses a diverse range of organisms, from free-living flagellates to parasitic protozoans^2^. This microorganism is globally distributed with its prevalence significantly varying among different populations. Notably, higher presence rates are reported in regions with lower socio-economic development, where prevalence can reach nearly 100%^3^. The role of *Blastocystis* in host health remains unclear. Historically considered an intestinal parasite, it has been linked to multiple gastrointestinal disorders, including irritable bowel syndrome and inflammatory bowel disease^4,5^. However, research findings regarding its role as a causative agent for these conditions are inconsistent. Some microbiome studies have identified *Blastocystis* as the most abundant eukaryotic commensal in the human gut^6–8^. Further research suggests that *Blastocystis* has a potentially beneficial role and is more prevalent in healthy individuals^1^. Specifically, human gut microbial communities that harbor *Blastocystis* have been associated with improved postprandial glucose responses, decreased body adiposity, favorable short-term cardiometabolic biomarkers, healthier dietary practices, and other positive health indicators^1,9^.

The discrepancies could arise from the significant genetic diversity of *Blastocystis* and its complicated links with gut microbiota. *Blastocystis* is currently categorized into at least 28 distinct subtypes (STs) with low host specificity^10^. Eight subtypes are identified in humans, with ST1, ST2, and ST3 being the most common^11^. In non-human primates (NHPs), both zoonotic (ST1–ST5, ST7, ST8, ST10), and non-zoonotic (ST11, ST13, ST15, and ST19) have been identified^6,12^. Notably, the co-occurrence of multiple subtypes within individuals (within-host diversity) is more common than previously thought for both animals and humans^13^. The pathogenicity of *Blastocystis* seems to be subtype-specific. For example, ST1 and ST4 can prevent DSS-induced colitis in a murine model, while ST7 exacerbates the disease^14,15^. In addition, emerging studies have shown that *Blastocystis* colonization influences gut microbiota composition, suggesting that its influence on health and disease may be closely linked with its interactions with gut bacteria (as reviewed in^6^). These interactions are complicated by the distinct biological traits exhibited by different *Blastocystis* subtypes, which include variations in growth rates and genetic repertoires, leading to both beneficial or adverse interactions with gut microbial communities. Indeed, for example, *Blastocystis* ST4 has been reported with beneficial effects on intestinal commensal bacteria and an inhibitory role on pathogenic *B. vulgatus*^16^, while ST7 exerts its pathogenic effects through disruption of the gut microbiota^17^. Thus, it has been proposed that microbial composition linked to *Blastocystis* likely depends on specific subtypes^6,7^. Despite this, investigations into the interactions between gut microbiota and different subtypes remain largely underexplored, especially regarding their co-occurrence patterns within the host.

In this study, we explored the relationships between the gut microbiota and *Blastocystis* intra-host diversity in a discovery cohort of *Macaca fascicularis* and a validation cohort of *Macaca mulatta* at subtype resolution. Unlike the previous studies involving clinical or community-dwelling human populations, our research here strictly excluded the effects of other common intestinal parasites and took advantage of the identical parameters of captive NHPs, including genetic backgrounds, living habits, environments, and diet, to control confounding effects. Through multifaceted approaches, including experiments on a new NHP cohort, our findings provide causal insights into how host gut microbiota relates to both the absolute and relative abundances of *Blastocystis* in the context of within-host heterogeneity.

## Results

### Cohort descriptions of non-human primates (NHPs)

To investigate the links between *Blastocystis* and the host gut microbiome, we collected fecal samples from captive *Macaca fascicularis* (discovery cohort, n = 348) and *Macaca mulatta* (validation cohort, n = 72) in Yuanjiang City, Yunnan Province, China (**Fig. 1a**). All primates in both cohorts shared the same diet and living environments. The presence of *Blastocystis* in the feces was diagnosed using a combination of PCR and qPCR methods targeting the specific *SSU* rRNA gene. We observed an unexpectedly high prevalence of *Blastocystis* at 97.9% (411 out of 420 samples), illustrating its widespread occurrence in these non-human primates (NHPs). To control for the influence of other prevalent intestinal parasites, we employed PCR-based methods to screen for various protozoans (including *Cryptosporidium* spp., *Giardia duodenalis*, *Enterocytozoon bieneusi*, *Cyclospora* spp., and Trichomonads) and common helminths (i.e., Nematodes and tapeworms). Samples with confirmed or suspected infections, or those that met the exclusion criteria were excluded from further analyses. Ultimately, we included 126 *Blastocystis* carriers (n = 100 for *M. fascicularis* and n = 26 for *M. mulatta*) for further microbiome analyses (**Fig. 1b**).

**Figure 1.**
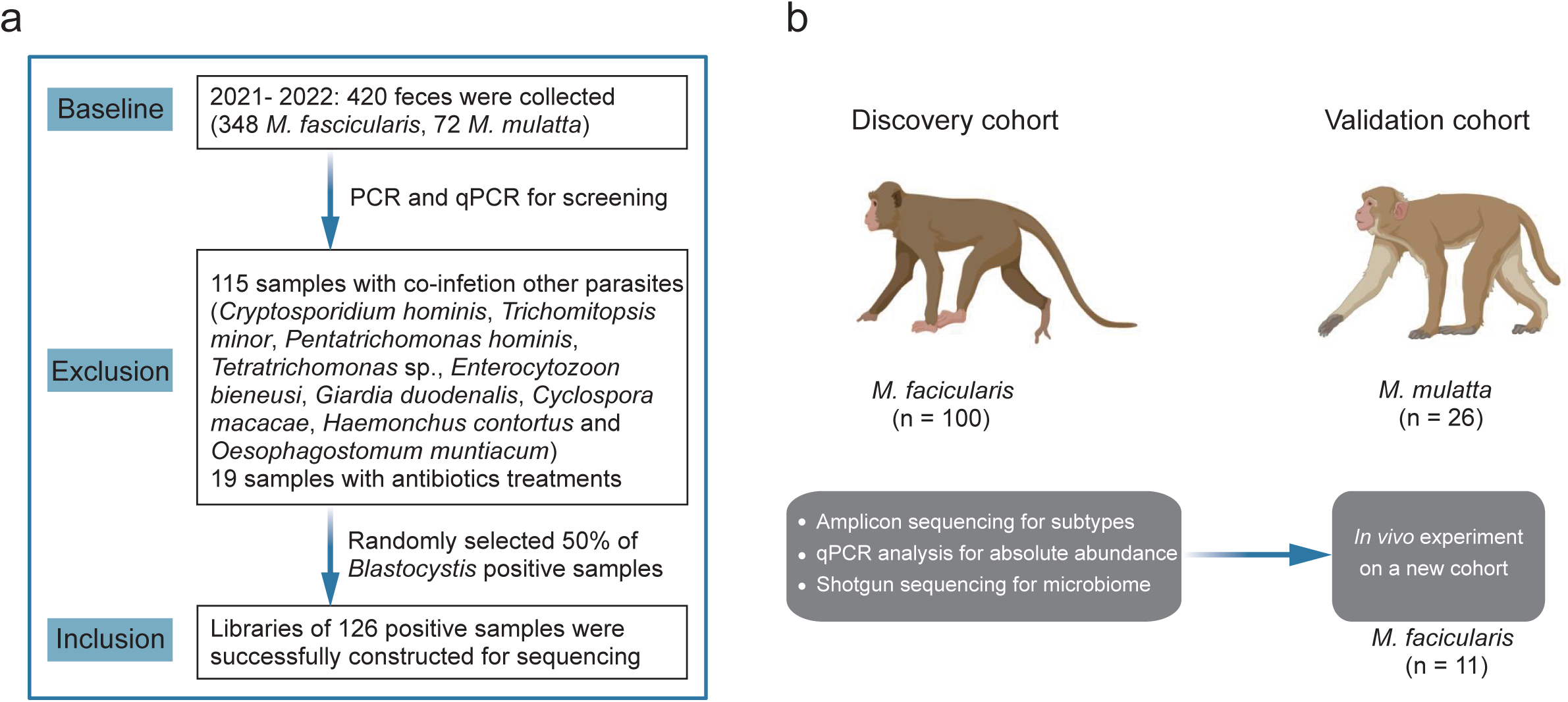
The overview of the study design. (**a**) Inclusion and exclusion of the samples in the study. (**b**) Overview of the study cohorts and methodologies employed. A discovery cohort of *M. facicularis*, a validation cohort of *M. mulatta*, and an independent experimental cohort of *M. facicularis* were used in this study. All the included samples were subject to a comprehensive analysis based on multifaceted techniques of PCR, qPCR, amplicon sequencing, and shotgun sequencing.

Each sample underwent microbiome profiling via metagenomic shotgun sequencing, yielding at least 40.1 million high-quality reads. We utilized amplicon sequencing of the *SSU* rRNA to estimate within-host *Blastocystis* genetic diversity in individual carriers, which produced a minimum of 42,126 reads per sample. In addition, we also determined the absolute abundance of *Blastocystis* in each sample using qPCR (**Fig. 1b**). Due to the identification of only 9 non-carrier controls (8 *M. fascicularis* and 1 *M. mulatta*), we did not include these samples in any further analysis to avoid potential statistical bias from the unequal sample sizes. With this comprehensive dataset, we tried to gain insights into the alterations of the host gut microbiome associated with *Blastocystis*.

### Identifiable within-host subtype diversity patterns of *Blastocystis* in non-human primates

Currently, at least 28 subtypes of *Blastocystis* have been reported^10^. Using the amplicon sequencing reads for *Blastocystis*, we identified 65 operational taxonomic units (OTUs) clustered at 97% identity across all samples, belonging to four subtypes: ST1, ST2, ST3, and ST5 (**Fig. 2a** and **Table S1**). Each subtype or sample exhibited high OTU diversity, with ST1 displaying up to 34 OTUs and individual samples containing as many as 39 OTUs (**Fig. S1**). This suggests a complex within-host genetic heterogeneity of *Blastocystis* in the gut of NHPs.

**Figure 2.**
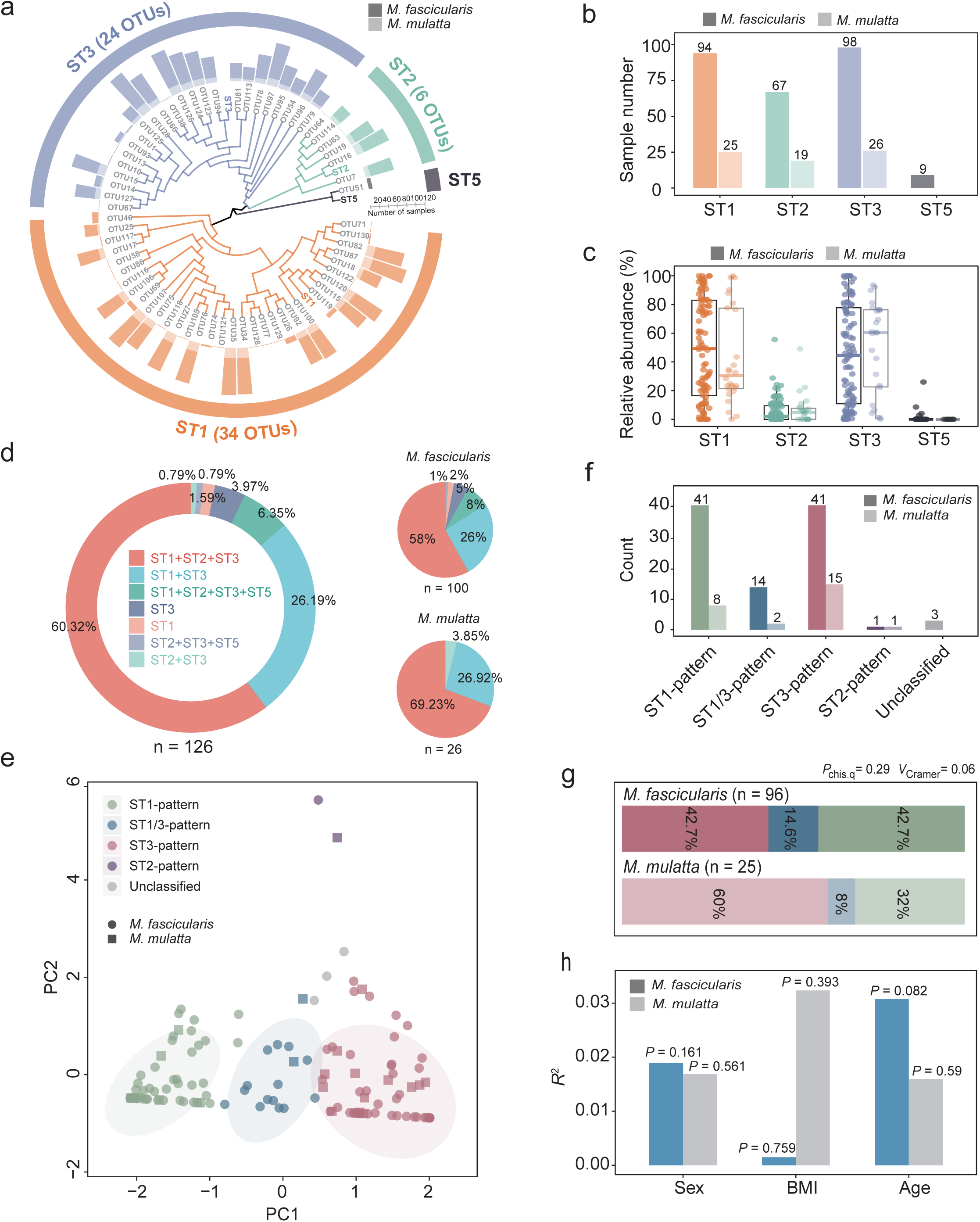
The characteristics of concurrent patterns of *Blastocystis* in NHPs. (**a**) Phylogenetic tree of *Blastocystis* subtype diversity at the OTU level. The OTUs with highlighted colors represent the reference sequence for each subtype. The Barplot indicates the number of samples with the presence of each subtype in amplicon sequencing. (**b**) The distribution of each *Blastocystis* subtype. (**c**) The relative abundance of each subtype in the samples. (**d**) The diversity of within-host subtype co-occurrence (concurrent patterns) of *Blastocystis* in NHPs. (**e**) The distribution of subtype concurrent patterns. The patterns were clustered using the fuzzy K-means (FKM)-based method. (**f**) The number of samples within each pattern. (**g**) The prevalence of the three dominant patterns (ST1-pattern, ST1/3-pattern, and ST3-pattern). (**h**) The effect size and significance of each host variable (sex, BMI, and age) on PCA of subtype concurrent patterns in PERMANOVA analysis. The *R*^2^ value represents the effect size for each variable. The *P* values were determined using the chi-squared test in panel g and using PERMANOVA analysis for panel h. The box plot represents the 25th percentile, median, and 75th percentile and whiskers stretch to 1.5 times the interquartile range from the corresponding hinge.

We compared the prevalence and relative abundance of each subtype across the samples. We found that ST1 and ST3 were the most prevalent subtypes, appearing in 94.44% (119/126) and 98.41% (124/126) of samples, respectively (**Fig. 2b and 2c**). Co-occurrence of these two subtypes with one another or with other subtypes within a host was notably common in this study. Specifically, the ST1+ST2+ST3 combination was the most prevalent, occurring in 60.32% of samples (76/126), followed by ST1+ST3 (26.19%, 33/126), ST1+ST2+ST3+ST5 (6.35%, 8/126), ST2+ST3 (0.79%, 1/126), and ST2+ST3+ST5 (0.79%, 1/126) (**Fig. 2d**). Similar trends were observed in both *M. fascicularis* and *M. mulatta* (**Fig. 2d**), highlighting a high within-host subtype diversity of *Blastocystis* in NHPs.

Based on the relative abundance of the two primary subtypes in each sample, we applied the fuzzy K-means (FKM)-based method to robustly classify the presence of *Blastocystis* in 121 NHPs into three main patterns (hereafter referred to as ‘**concurrent pattern**’): ST1-pattern, ST3-pattern, and ST1/3-pattern (**Fig. 2e** and **Fig. S2**). These mixed subtype patterns are characterized by the dominance of either ST1, ST3, or a balanced presence of both, respectively. Notably, the ST1-pattern and ST3-pattern predominated over the ST1/3-pattern in terms of prevalence (**Fig. 2f**) within both *M. fascicularis* and *M. mulatta*, suggesting competitive interactions between subtypes within the *Blastocystis* populations (**Fig. 2g**). The PERMANOVA analysis indicated that differences in host parameters, including age, BMI, or sex among the NHPs could not adequately explain the clustering of observed patterns in NHPs (**Fig. 2h**). These findings support the notion that the presence of *Blastocystis* can be characterized by subtype concurrent patterns in the carriers of both *M. fascicularis* and *M. mulatta*.

### Host gut microbiome variations characterize subtype concurrent patterns

Next, we examined the associations between the concurrent subtype patterns of *Blastocystis* and gut microbial compositions. To streamline our analysis, we only present the findings for the discovery cohort, *M. fascicularis*, and will specify the key findings for the validation cohort, *M. mulatta* in the end. The Shannon index (**Fig. 3a**) indicated that the level of alpha diversity was comparable across concurrent patterns. We assessed compositional variations using Principal Coordinates Analysis (PCoA) based on Bray-Curtis distance at the species level and found a significant shift in the gut microbiome among *Blastocystis* subtype presence patterns (**Fig. 3b**). Furthermore, our PERMANOVA analysis revealed that the concurrent patterns accounted for more variations in gut microbiota than known host covariates such as age, BMI, and sex (**Fig. 3c**). As illustrated by the Bray-Curtis distance between samples (**Fig. 3d**), gut microbial compositions in ST1-pattern samples were structurally more similar to those in ST1/3-pattern samples than to those in ST3-pattern samples, indicating microbiota compositional differences across the samples with different co-occurrence of subtypes.

**Figure 3.**
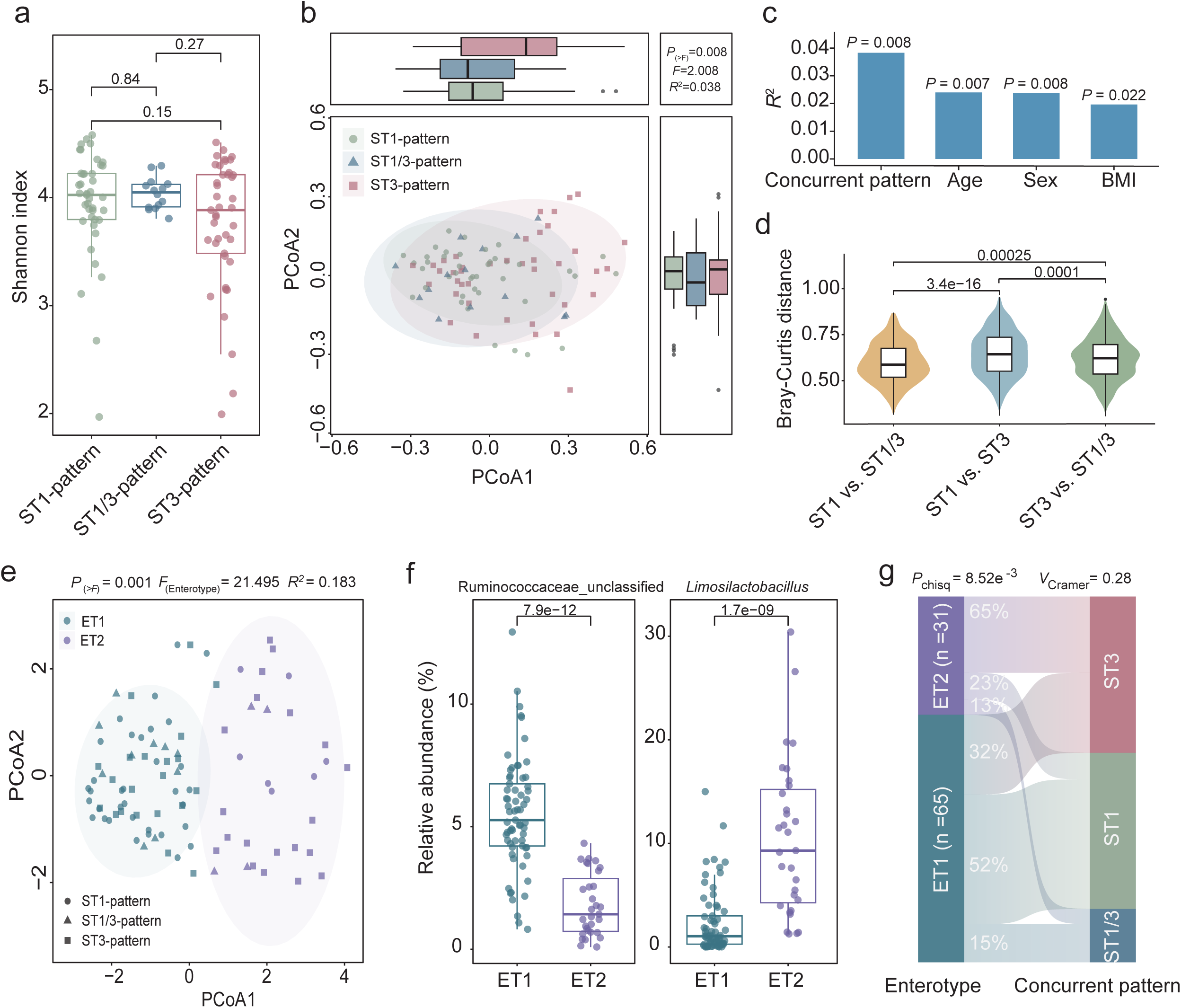
The gut microbiome across subtype concurrent patterns in *M. fascicularis*. **(a)** The Shannon index across subtype concurrent patterns. **(b)** The Principal Coordinates Analysis (PCoA) of the gut microbiota at the species level based on Bray-Curtis distances. **(c)** Effect sizes of concurrent patterns and sample covariates on the PCoA analysis. The *R*^2^ value for each variable was determined using the PERMANOVA analysis. **(d)** The Bray-Curtis distances for inter-group comparisons. **(e)** The gut microbiome enterotypes in subtype concurrent patterns. Enterotypes were identified using Jensen-Shannon distance (JSD) and partitioning around medoid (PAM) clustering at the genus level. **(f)** Relative abundances of Ruminococcaceae and *Limosilactobacillus* within the enterotypes. **(g)** Associations between enterotypes (ET1 and ET2) and subtype concurrent patterns. The number of samples is shown in each panel. The confounding variables (sex, age, and BMI) were adjusted in the analysis for panels b and e. The *P* values were determined by the Wilcoxon rank-sum test for panels a, d, and f, by the PERMANOVA test for b, c, and e, and by the chi-squared test for panel g. The box plot represents the 25th percentile, median, and 75th percentile and whiskers stretch to 1.5 times the interquartile range from the corresponding hinge.

Furthermore, we examined microbial variations associated with the concurrent patterns in terms of enterotype, which provides a structurally broader understanding of the relationships involved. Using JSD-PAM-based methods, we classified the gut microbiota of *M. fascicularis* into two distinct enterotypes, characterized by the dominant genera: an unclassified genus in Ruminococcaceae (ET1) and *Limosilactobacillus* (ET2) (**Fig. 3e and 3f**). PCoA analysis, based on Jensen-Shannon divergence, demonstrated a significant distinction between samples from ET1 and ET2. The enterotype analysis clearly illustrated distinct associations with microbial structures across different concurrent patterns (**Fig. 3g**). Specifically, ET1 was enriched in both the ST1-pattern and ST1/3-pattern, whereas ET2 was predominantly identified in the ST3-pattern (**Fig. 3g**). Collectively, these findings suggest that the subtype concurrent patterns of *Blastocystis* are correlated with compositional and structural variations in the gut microbiome.

### *Blastocystis* load explains the associations between concurrent patterns and gut microbiota

To gain insights into the association between concurrent patterns and the gut microbiome in *M. fascicularis*, we examined factors related to the co-occurrence of subtypes, specifically the abundance of each subtype. Our analysis revealed that the total *Blastocystis* load in each sample exhibited the strongest explanatory power for variations in gut microbiota (**Fig. 4a**). This finding suggests that the absolute abundance of *Blastocystis* in each sample represents a critical feature in differentiating concurrent patterns. Indeed, qPCR analysis indicated that samples exhibiting the ST1 or ST1/ST3 patterns contained significantly higher amounts of *Blastocystis* compared to those with the ST3-pattern (**Fig. 4b**). Notably, we found a significant positive correlation between the relative abundance of the ST1 subtype and *Blastocystis* load across samples, while the relative abundance of ST3 demonstrated a negative correlation with *Blastocystis* levels (**Fig. 4c**). This suggests that the dominant subtype within each concurrent pattern influences the overall quantity of *Blastocystis* present.

**Figure 4.**
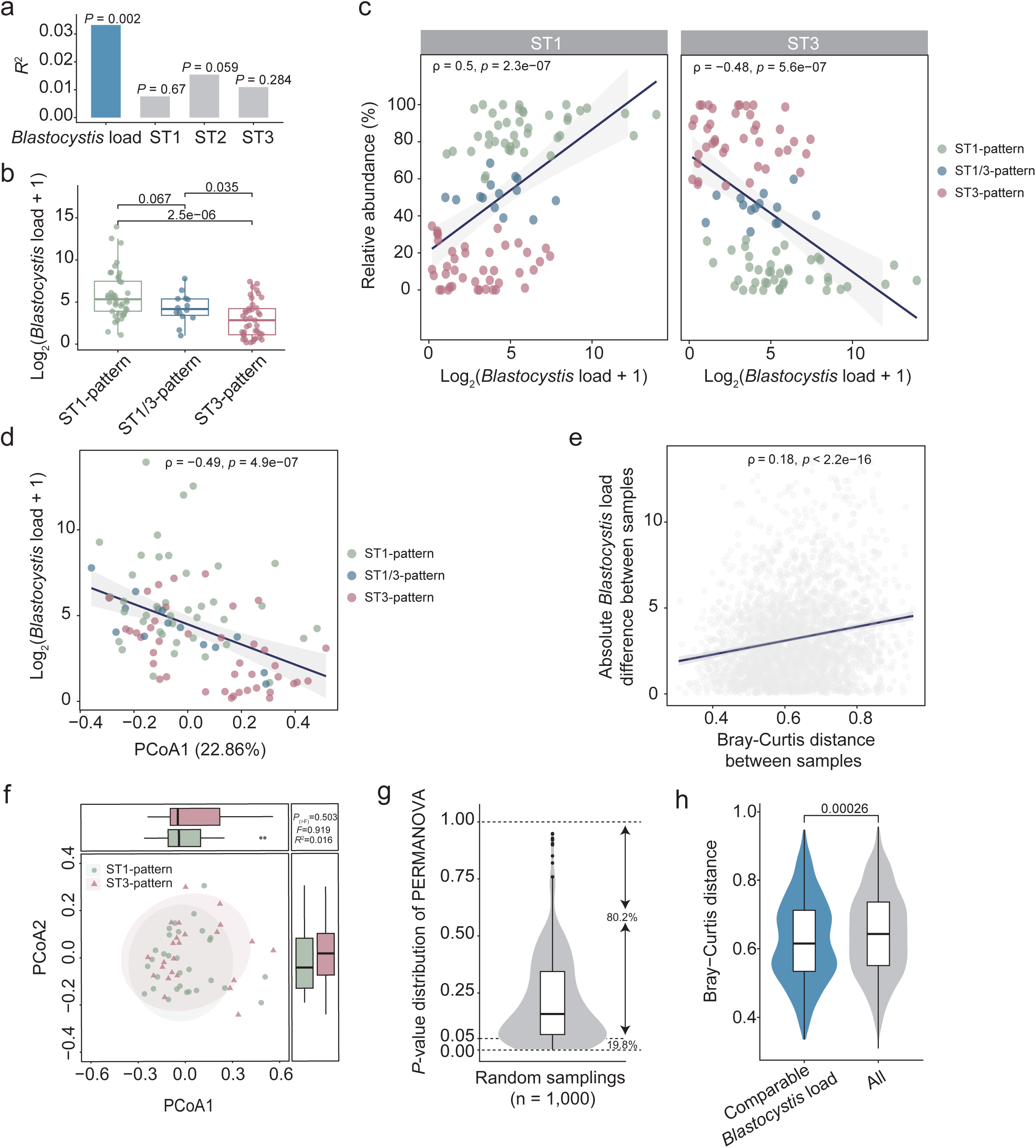
*Blastocystis* load explains the associations between subtype concurrent patterns and the gut microbiota in *M. fascicularis*. (a) Effect sizes of the absolute abundance of *Blastocystis* and relative abundance of each subtype related to concurrent patterns on the PCoA analysis of the gut microbiota (Fig. 3b) in the PERMANOVA analysis. **(b)** The absolute abundance of *Blastocystis* in each concurrent pattern. Log2-transformed values are shown for the Y-axis. **(c)** Spearman correlation between the relative abundance of ST1 (or ST3) and the total *Blastocystis* load in the sample. **(d)** Spearman correlation between *Blastocystis* load and the first principal coordinate (PCoA1) in PCoA analysis (Fig. 3b). **(e)** Spearman correlation between Bray-Curtis distance and *Blastocystis* load. **(f)** Bray-Curtis distance-based PCoA analysis for samples of ST1-pattern and ST3-pattern with the comparable *Blastocystis* load between them. **(g)** Distribution of *P* values from the PERMANOVA analysis by BootStrap sampling (n = 1,000). **(h)** Bray-Curtis distances between ST1-pattern and ST3-pattern for samples with comparable absolute abundance. The confounding variables (sex, age, and BMI) were adjusted in the analysis for panel f. The *P* values were determined by the PERMANOVA test for panels a and f, by the Wilcoxon rank-sum test for panels b and h, and by Spearman’s rank correlation coefficient test for panels c, d, and e. The box plot represents the 25th percentile, median, and 75th percentile and whiskers stretch to 1.5 times the interquartile range from the corresponding hinge.

Thus, our data suggest that the absolute abundance of *Blastocystis* is a key factor explaining the associations between gut microbial variations and concurrent patterns. Supporting this idea, we observed a significant correlation between the first principal coordinate (PCoA1) in the PCoA analysis (**Fig. 3b**) and the measured *Blastocystis* count in feces (**Fig. 4d**). Additionally, the Bray-Curtis dissimilarities in gut microbiomes between samples were significantly correlated with variations in absolute abundance of *Blastocystis* (**Fig. 4e**). To further investigate whether *Blastocystis* load dominates the associations between gut microbiome and concurrent patterns, we randomly selected ST1-pattern samples with a *Blastocystis* load comparable to that of ST3-pattern samples and compared their microbiome structures using PERMANOVA through a bootstrap resampling method (n = 1,000). PCoA analysis indicated that microbial structural differences diminished between the two patterns (**Fig. 4f and 4g**, PERMANOVA, *P* > 0.05, Bootstrap = 80.2%), accompanied by a significant reduction in Bray-Curtis distance (**Fig. 4h**). Collectively, these findings support the idea that *Blastocystis* load is a critical determinant in elucidating the associations between concurrent patterns and gut microbiota.

### Lactic acid bacteria reduce the absolute abundance of *Blastocystis* in carriers

Our above results suggest a close relationship between host gut microbiota and the abundance of *Blastocystis*; we thus investigated the microbial factors influencing *Blastocystis* load in carriers. We assessed the association between *Blastocystis* load and gut microbial compositions in *M. fascicularis* while controlling for other host covariates (BMI, sex, and age) using MaAsLin2. At the species level, we identified 92 bacteria that demonstrated a strong co-occurrence with *Blastocystis* (**Table S2** and **Fig. 5a**). Notably, species from the lactic acid bacteria group (*Limosilactobacillus* and *Lactobacillus*) ranked among the most enriched taxa that were negatively associated with *Blastocystis* load (**Table S2**).

**Figure 5.**
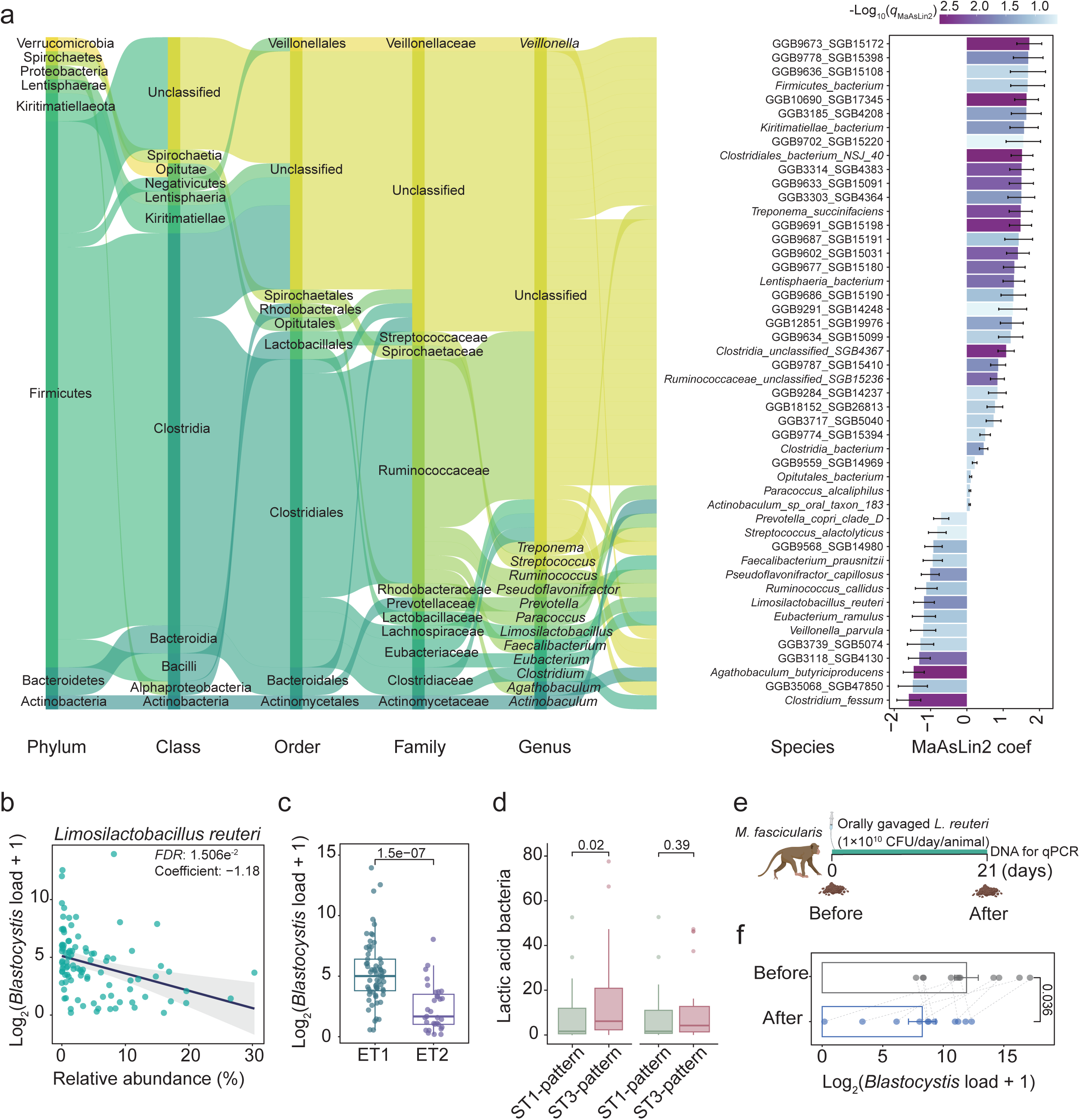
Impact of lactic acid bacteria on *Blastocystis* load in *M. fascicularis*. **(a)**The associations of microbial taxa with *Blastocystis* load in MaAsLin2 analysis. The top taxa ranked by q-value (q < 0.1) are shown at different levels. **(b)** Correlation between the relative abundance of *Limosilactobacillus reuteri* and *Blastocystis* load in MaAsLin2 analysis. **(c)** Comparison of *Blastocystis* load between samples of enterotypes (ET1 and ET2). **(d)** Relative abundance of lactic acid bacteria (*Lactobacillus* and *Limosilactobacillus*) in ST1- and ST3-patterns for the samples with comparable *Blastocystis* load. **(e)** Experimental design for an independent cohort of *M. fascicularis*. Oral administration of *L. reuteri* was performed daily. **(f)** Changes of *Blastocystis* load before and post the experiment. qPCR was used to examine the *Blastocystis* abundance in feces. The confounding variables (sex, age, and BMI) were adjusted in the analysis for panels a and b using MaAsLin2. The *P* values or *FDR* were calculated using MaAsLin2 for panels a and b, the Wilcoxon rank-sum test for panels c and b, and the Wilcoxon signed-rank test for panel f. The box plot represents the 25th percentile, median, and 75th percentile and whiskers stretch to 1.5 times the interquartile range from the corresponding hinge. The data of mean ± SD is shown for panels a and f.

For instance, *Limosilactobacillus ruteri* emerged as the predominant taxon (**Fig. 5b**). This aligns with our observation that ET2, enriched in *Limosilactobacillus*, exhibited a significantly lower *Blastocystis* load compared to enterotype ET1, which is enriched in Ruminococcaceae (**Fig. 5c**). Moreover, we found the level of lactic acid bacteria in the ST1-pattern was significantly lower than that in the ST3-pattern, whereas this trend disappeared if samples with comparable *Blastocystis* load were randomly selected for comparison (**Fig. 5d**).

These findings support the hypothesis that lactic acid bacteria could impact *Blastocystis* levels in primates. Indeed, a previous study has indicated that lactic acid bacteria may inhibit the proliferation of *Blastocystis in vitro*^18^. To test this hypothesis in NHPs *in vivo*, we recruited a new cohort of *Blastocystis* carriers of *M. fascicularis* (n = 11) and administered an *L. reuteri* strain (1x10^10^ CFU/day/animal) orally. We measured the absolute level of *Blastocystis* in their feces using qPCR (**Fig. 5e**). After 21 days of *L. reuteri* gavage, we observed a significant reduction in *Blastocystis* number compared to the pre-treatment levels (**Fig. 5f**). Collectively, these data suggest that lactic acid bacteria significantly influence *Blastocystis* load.

### The relationship between *Blastocystis* load and the gut microbiota is a common and predictive feature in NHPs

Our findings indicate that the gut microbial compositions are closely linked to the abundance of *Blastocystis* in *M. fascicularis*. This observation raised questions about whether this association is a common feature that could serve as an indicator of *Blastocystis* levels in non-human primates (NHPs). To explore this hypothesis, we first replicated our analyses in the cohort of *M. mulatta*. Using the same analytical methods, we identified comparable trends to those seen in *M. fascicularis*, confirming that *Blastocystis* load significantly correlates with microbial variations across different concurrent patterns (**Fig. 6a-c**). This conclusion is supported by extensive analyses of compositional and structural configurations, particularly regarding diversity and enterotypes (**Fig. 6a-c** and **Fig. S3**). Consistently, we found that the *Blastocystis* load was significantly lower in gut microbiota configurations characterized by a high abundance of *Limosilactobacillus* (**Fig. 6d**). Furthermore, we developed a machine-learning classifier utilizing the Random Forest (RF) method, partitioning the samples from both *M. fascicularis* and *M. mulatta* into discovery and testing subsets. The microbial biomarkers identified (**Fig. 6e**) were able to effectively predict *Blastocystis* load in both testing and validation datasets, achieving a high coefficient of determination (*R*^2^) (**Fig. 6f**). These results suggest that the microbial features associated with *Blastocystis* load are consistent across the two NHP species. Collectively, this data supports the hypothesis that the gut microbiome plays a predictive role regarding *Blastocystis* load in its carriers.

**Figure 6.**
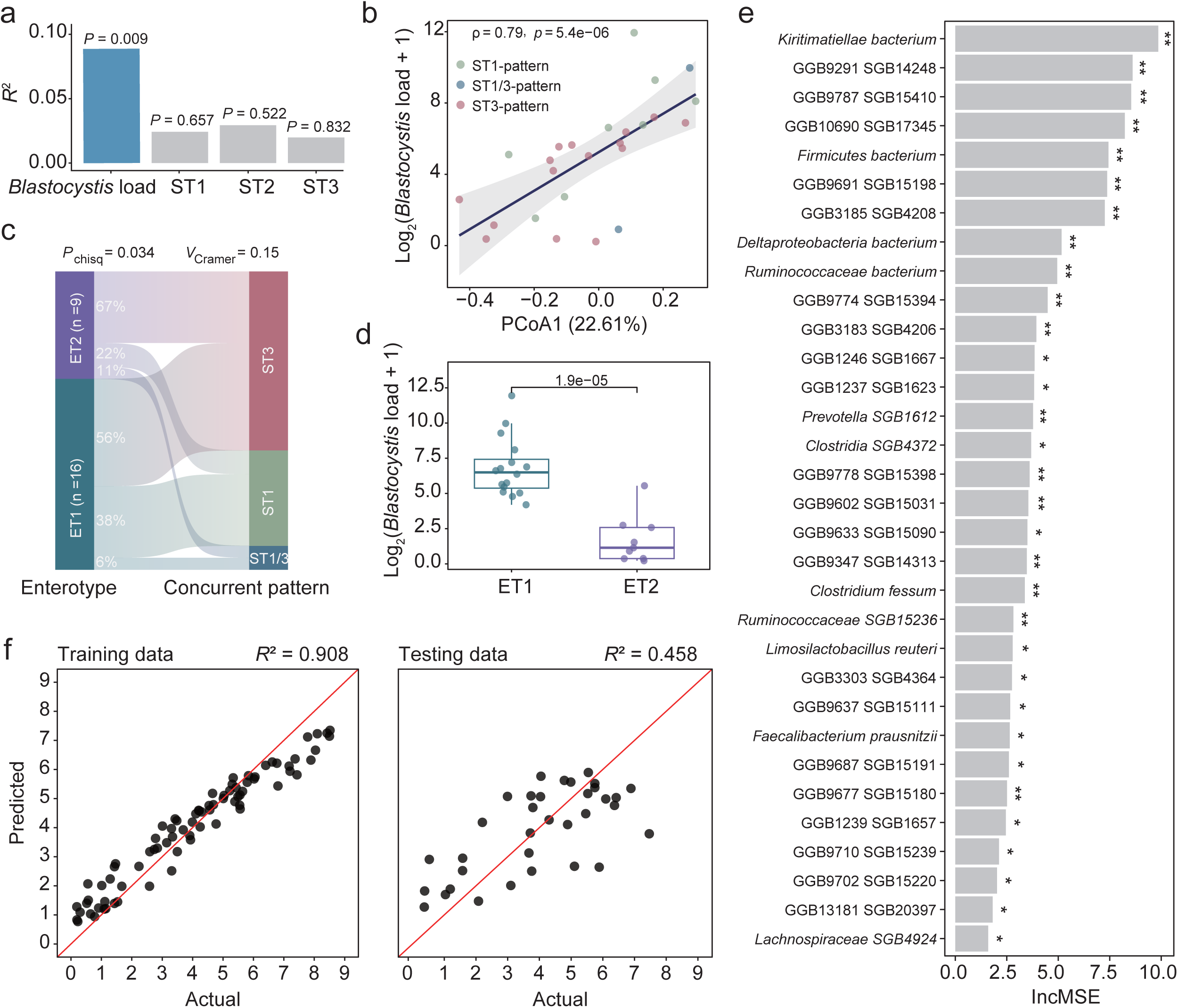
The gut microbiota composition can predict *Blastocystis* load in NHPs. (**a**) Effect sizes of each variable on the PCoA analysis of the gut microbiota for *M. mulatta.* PERMANOVA analysis was used to determine each variable’s effect size and significance. (**b**) Spearman correlation between *Blastocystis* load and PCoA1 of the microbial analysis in *M. mulatta*. (**c**) Associations between enterotypes (ET1 and ET2) and concurrent patterns (ST1/3, ST1, and ST3) in *M. mulatta*. (**d**) *Blastocystis* load in the samples with ET1 and ET2 in *M. mulatta*. (**e**) The identified markers (n = 32) in Random Forest (RF) analysis for the discovery cohort dataset (n = 79). These markers are ranked based on their percentage increase in mean squared error (% IncMSE). (**f**) The performance of the RF model in predicting training or testing datasets. The *P* values were determined using PERMANOVA analysis for panel a, Spearman’s rank correlation coefficient test for panel b, and the Wilcoxon rank-sum test for panel d. \**P* value < 0.05; \*\**P* value < 0.01.

## Discussions

*Blastocystis* is a widespread protist found in both animals and humans globally. The notably high prevalence of *Blastocystis* in non-human primates, specifically the 97.9% presence rate recorded in our study, highlights its ecological significance within primate gut microbiomes and reinforces the notion that even under controlled conditions with captive *M. fascicularis* and *M. mulatta*, *Blastocystis* continues to be a dominant component of the microbial community. Thus, understanding the relationships between *Blastocystis* and the other parts within the host gut microbiome is of high importance. Given the substantial inter-subtype genetic variation, characterizing these subtypes is essential for understanding the connections between microbiota composition and *Blastocystis*. Accordingly, controlling for environmental variability (e.g., diet, habit, and genetics), the current study examines the relationship between gut microbiota and *Blastocystis* while considering its within-host diversity.

To the best of our knowledge, this study is the first cohort investigation focusing on the impact of within-host genetic diversity of *Blastocystis* on the gut microbiota. Our findings highlight the significant prevalence of mixed-subtypes existence within a single host in non-human primates. Although often underexplored, previous studies indicate that the presence of mixed subtypes within a specimen is more common than previously recognized^13,19–21^. For example, co-occurrence involving multiple *Blastocystis* subtypes was identified in 62.5% of positive samples from birds^22^ and 22% of human cases^13^. In our study, employing an amplicon sequencing method^20^, we discovered that the co-occurrence of multiple subtypes accounts for over 90% of the total carriers involved. Additionally, our results revealed a high intra-subtype variability and a substantial number of sequence variants even at the operational taxonomic unit (OTU) level. Given that our findings, along with previous research, consistently demonstrate a strong correlation between subtypes and microbial variation, it is crucial to consider intra-subtype and even intra-isolate genetic heterogeneity in *Blastocystis*-related microbiome studies.

Our findings underscore the critical role of absolute abundance in examining the interplay between *Blastocystis* and gut microbiota. Numerous studies have reported subtype-dependent associations with gut microbiota, with different subtypes exhibiting varying effects in both human and murine models^6,7^. For instance, the presence of ST4 was correlated with elevated levels of *Sporolactobacillus* and *Candidatus carsonella* in Swedish travelers, while ST3 did not demonstrate such significant relationships^23^. Additionally, an inverse correlation between ST3 and ST4 and *Akkermansia* abundance was observed in fecal samples from a distinct cohort study^7^. These findings highlight the complex nature of *Blastocystis*, as different subtypes possess divergent biological characteristics, including growth rates, colonization niches, and host preferences^17,24^. However, the underlying mechanisms remain largely unclear. In this study, we dissected the factors associated with microbial variations between concurrent patterns of subtypes, emphasizing that the absolute abundance of *Blastocystis* is a key determinant for understanding its associations with gut microbiota. This may reflect the biological variations, particularly in growth rates or colonization dynamics among subtypes. Supporting this idea, our results revealed a positive correlation with ST1 and a negative correlation with ST3 regarding absolute abundance in fecal samples. This is in line with findings in a cohort study of human populations, which indicated a higher load in ST1 carriers compared to ST3 carriers^7^, as well as an *in vitro* study that demonstrated a higher growth rate for ST1 than for ST3 when cultured in DMEM medium^25^.

Our findings provide preliminary causal insights into the relationship between the absolute abundance of *Blastocystis* and the gut microbiota. Notably, our results suggest that this relationship cannot be solely attributed to the amount-dependent effects of *Blastocystis*. Instead, our data indicate that the load of *Blastocystis* may be influenced, at least in part, by the host’s microbial composition, particularly the presence of lactic acid bacteria. Our *in vivo* experiment demonstrated a strong inhibitory effect of lactic acid bacteria on *Blastocystis* levels in non-human primates. This aligns with previous research, which found that *in vitro* co-incubation with either *L. rhamnosus, L. lactis, or E. faecium* inhibited the growth of *Blastocystis*^18^. These findings strongly suggest that a higher abundance of these beneficial bacteria may enhance a host’s resistance to *Blastocystis* colonization, although the precise underlying mechanisms require further investigation. Furthermore, it is reasonably speculated that the presence of *Blastocystis* in primates can lead to alterations in microbial composition and diversity, as reported in various studies on different models. Nonetheless, this research underscores the potential of using *L. ruteri* as a prophylactic approach to regulate *Blastocystis* colonization.

In summary, this study underscores the ecological significance of *Blastocystis* in primate guts and emphasizes the need to consider subtype diversity and absolute abundance in microbiome studies. The inhibitory role of lactic acid bacteria opens avenues for probiotic interventions in non-human primates. Future research should explore mechanistic pathways and validate findings in human cohorts and wild populations.

## Methods

### Ethics declarations

All animal work was conducted according to the guidelines of the Kunming Biomed International (KBI) Animal Experiment Management and Ethics Committee (No. KBI K001123083-01, 01).

### Study cohorts and fecal sample collection

Fecal samples were collected from captive *Macaca fascicularis* and *Macaca mulatta* between 2021 and 2022 in Yuanjiang City, Yunnan Province, China. The monkeys were individually housed in separate cages. Data on their age, sex, BMI, diet, and medication history were recorded. Fresh samples (5-10 g/animal) were collected and immediately transferred into a sterile collection tube containing DNA preservation solution (Phygene, Cat#PH1408). The samples (n = 348 for *M. fascicularis* and n = 72 for *M. mulatta*) were aliquoted and stored at -80°C until use. Stool DNA was extracted using E.Z.N.A® Stool DNA Kit (OMEGA, Cat#D4015-02) according to the manufacturer’s instructions.

### Inclusion and exclusion criteria

To identify *Blastocystis*, a combination of PCR-based and qPCR-based methods was used. First, primers targeting the small subunit rRNA (*SSU* rRNA) gene (BhRDr: 5′-GAGCTTTTTAACTGCAACAACG-3′ and RD5: 5′-ATCTGGTTGATCCTGCCAGT-3′)^26^ were used for screening. Second, for *Blastocystis*-negative samples, real-time quantitative PCR (qPCR) was used for verification^27^ (See below). Sanger sequencing on the *SSU* rRNA region was used to confirm the presence of *Blastocystis* in the positive samples in either method.

For exclusion, the common protozoans (*Cryptosporidium* spp.^28^, *Giardia duodenalis*^29,30^, *Enterocytozoon bieneusi*^31^, *Cyclospora* spp.^32^, and Trichomonads^33^) as well as helminths (using universal primers)^34,35^ were screened using PCR methods. Nine parasite species (*Cryptosporidium hominis*, *Trichomitopsis minor*, *Pentatrichomonas hominis*, *Tetratrichomonas* sp., *Enterocytozoon bieneusi*, *Cyclospora macacae*, *Giardia duodenalis*, *Haemonchus contortus*, *Oesophagostomum muntiacum*) were found in the samples after the screening. The samples containing these pathogens were excluded. In addition, NHPs with active gastrointestinal diseases (e.g., diarrhea and vomiting), active chronic viral infections, or a history of antibiotics or probiotics use within the past 3 months were also excluded from the further study.

### Real-time quantitative PCR (qPCR) measurement

For qPCR analysis, the absolute number of *Blastocystis* cells in feces was estimated according to the previously published protocol^27^. Briefly, 2 μL of extracted DNA was mixed with 10 μL of 2× GoTaq® qPCR Master Mix (Promega, Cat#A6002), 1 μL (0.5 μM) of each primer (BL18SPPF1: 5′-AGTAGTCATACGCTCGTCTCAAA-3′ and BL18SR2PP: 5′-TCTTCGTTACCCGTTACTGC-3′), and 6 μL of nuclease-free water, for a total reaction volume of 20 μL. The qPCR was performed on an Applied Biosystems (ABI) 7500 Real-Time PCR system (Thermo Fisher Scientific) with an initial denaturation step at 95 °C for 2 min, followed by 45 cycles of 95 °C for 15 s and 68°C for 1 min. Standard curves were established using genomic DNA from an *in vitro* culture of *Blastocystis* (ST1), according to the previously reported protocol^27^. The number of *Blastocystis* cells per milligram of feces was then calculated based on the comparison of CT values in each sample against the standard curve.

### Amplicon sequencing and analysis

For estimating the within-host diversity of *Blastocystis* subtypes, an amplicon sequencing strategy was employed, using a protocol as previously described^20^. In brief, 126 *Blastocystis-*positive samples were amplified using the PCR method with primers targeting a specific region of the *SSU* rRNA gene (Blast505_532F and Blast998_1017R)^36^ and linking with Illumina overhang adapter sequences at the 5′ end. The final libraries were quantified using Qubit fluorometric quantitation (Thermo Fisher Scientific) and were sequenced on the NextSeq 2000 platform (Illumina), following the manufacturer’s recommendations. The paired-end reads were processed and analyzed using a pipeline that incorporates fastp (v0.19.6)^37^, FLASH (v1.2.11)^38^, and UPARSE (v11)^39^. The raw reads were processed using fastp to filter low-quality reads (--cut_window_size = 50, --cut_mean_quality = 20, --length_required = 50, --n_base_limit = 1). The filtered reads from each sample were merged into raw tags using FLASH (v1.2.11) (--min-overlap = 10). UPARSE was used to cluster the OTUs at 97% similarity and remove chimeras. The resulting OTUs were then searched by BLASTn against the NCBI database for species classification (e-value <1e^-5^). IQTree2^40^ was applied to construct a bootstraps maximum-likelihood tree with parameters “-m MFP -bb 1000” using the representative sequences of OTU.

### Construction of concurrent patterns of *Blastocystis*

We used a fuzzy K-means (FKM)-based method^41^ to identify a distinct set of concurrent patterns of *Blastocystis* subtypes strongly supported by the relative abundance of each subtype in samples. This method employs the FKM algorithm to initially cluster the samples into preliminary groups, followed by further filterings. The pairwise distance matrix used in the FKM clustering is computed using the formula:

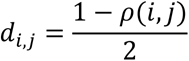

Where 𝑑_𝑖,𝑗_ and 𝜌(𝑖, 𝑗) represent the distance and Pearson correlation between samples i and j, respectively. The minimum membership threshold for sample assignment is set to 0.8. If a sample exhibits membership in multiple groups, it is assigned to the group with the highest membership value. If a sample’s membership values for all groups fall below the threshold, it is classified as “unclassified”.

The R function ‘fanny’ from the R package cluster (v2.1.6) was used for fuzzy K-means clusters. We tested the number of clusters (k) ranging from 1 to 7, and the optimal k was determined as the one that clustered the largest number of samples (k = 4).

### Metagenomic sequencing and analyses

DNA libraries were prepared using the NEXTFLEX Rapid DNA-Seq Kit (Bioo Scientific), following the manufacturer’s recommendations. Sequencing was performed on the NovaSeq 6000 platform (Illumina). The raw shotgun sequencing data were analyzed using the bioBakery metabolomics pipeline to generate both taxonomic and functional profiles. The KneadData (v.0.10) (https://github.com/biobakery/kneaddata) was used to detect and remove reads derived from the hosts (*M. fascicularis*, genome version: GCF_012559485.2; *M. mulatta*, genome version: GCF_003339765.1). The adapter sequences at 3’ and 5’ ends of reads were mass-cut and the reads with a length less than 50 bp, an average base mass value less than 20, or having ‘N’-base were discarded. Taxonomic identification were performed using MetaPhlan4^42^ (v.4.0.3) using the standard reference database (mpa_vJan21_CHOCOPhlAnSGB_202103) with default parameters.

### Enterotype analyses

For enterotype analysis, the gut bacterial composition was clustered at the genus level across all samples using a previously described method^43^. To minimize noise, bacterial taxa detected in fewer than 20% of the samples were excluded from the analysis. Clustering was conducted using partitioning around medoids (PAM) based on the Jensen-Shannon distance (JSD) between samples. The optimal number of clusters was determined using the Calinski-Harabasz index. To validate the clustering results and identify dominant bacterial taxa within each cluster, Between-Class Analysis (BCA) was applied.

### Random Forest analysis

We implemented the Random Forest (RF) regression model using the R package randomForest (v4.7-1.1) to estimate the relationship between the relative abundance of microbiota and the absolute abundance of *Blastocystis*. To reduce the impact of outlier values, we used samples with a *Blastocystis* load of less than 512 (n = 118). The original dataset with the samples from *M. fascicularis* and *M. mulatta* was randomly split into a training (discovery) set and a validation set, with a 7:3 ratio. The optimal markers to predict the absolute abundance of *Blastocystis* for the training dataset were identified using the Boruta algorithm, implemented in the R package Boruta (v8.0.0). The statistical significance (*P* value) for each marker was assessed using the R package rfPermute (v2.5.2) (https://github.com/EricArcher/rfPermute). Only markers with a *P* value < 0.05 in the permutation test were kept and then ranked by decreased %IncMSE (% Increase in Mean Squared Error). The final “bagged” RF regression model, based on the optimal set of identified microbial features, was subsequently applied to evaluate predictive efficiency in both the training and validation datasets. The performance of the regression model was assessed using the coefficient of determination (*R*^2^), which was calculated by the formula:

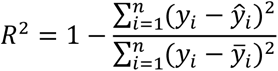

𝑦_𝑖_ and *ŷ_i_* represent Indicates the actual values and predicted value of absolute abundance of *Blastocystis* in sample i. *ȳ_i_* represent the mean value of all observations (𝑦_1_, 𝑦_2_, 𝑦_3_, …, 𝑦_𝑖_).

### *L. reuteri* intervention experiment

The *L. reuteri* A21041 strain was kindly provided by Guangxi Key Laboratory of Longevity Science and Technology, Nanning. The lyophilized powder of this strain was dissolved in sterile PBS (Gibco, Cat#C20012500BT) and cultured on an MRS solid medium plate for 24 h at 37°C in anaerobic conditions to estimate the number of viable bacteria. We recruited a new group of *Blastocystis* carriers (*M. fascicularis*, n = 11) and housed them in separate cages at an ambient temperature of 22 ± 1°C and a 12/12 h light/dark cycle. The lyophilized powder of the *L. reuteri* strain was thoroughly mixed with normal saline (Servicebio, Cat#G4702) and administered via oral gavage (1x10^10^ CFU/day/animal). All *M. fascicularis* received the *L. reuteri* intervention for 21 consecutive days. Fecal samples were collected every 24 h before the intervention and qPCR was used to evaluate *Blastocystis* load in each sample.

### Statistical analysis

Statistical analyses were performed in R (v4.4.0). The ‘vegan’ package (v2.6-8) was used for α-diversity analysis. The MaAsLin2 (v1.18)^44^ package was used for controlling for covariate adjustment (sex, age, and BMI) for both comparison and association analyses. β-diversity was measured based on the Bray-Curtis distance, with statistical significance assessed via Permutational Multivariate Analysis of Variance (PERMANOVA) using the ‘adonis2’ function in the vegan package (v2.6-8). The aPCoA package (v1.3)^45^ was also used to adjust for covariates (sex, age, and BMI) in PCoA or PCA analysis. Spearman’s rank correlation coefficient was computed using the ‘cor.test’ function, and differences between groups were assessed using the Wilcoxon rank-sum test (two-sided, confidence level = 0.95). Benjamini-Hochberg procedure (*FDR*) was used to correct *P* values for multiple hypothesis testing.

## Acknowledgments

This work was supported by the Guangxi Key Laboratory of Longevity Science and Technology (Open Fund Project No. gxkllst-20241001), and the Gansu Provincial Natural Science Foundation (23JRRA1514). We thank the workers from Kunming Biomed International for collecting samples and performing experiments on monkeys.

## Data Availability

Data are available in the main text or the supplementary materials. Sequences have been deposited to NCBI with accession number: PRJNA1222911. Original R scripts, metadata, and OTU tables are available on GitHub at (https://github.com/Axolotl233/Blastocystis_microbes).

## Author contributions

SW and WJM designed the study. WJM, PPM, YHL, YZ, YGW, LXC, YYP, WFL, HP, and LZ collected samples and performed experiments. WJM and PPM analyzed the microbiome data and prepared figures. SW, WJM, and PPM prepared the manuscript with contributions from all co-authors.

## Supplementary Figure legends

**Figure S1. The number of OTUs in individual samples of NHPs.**

**Figure S2. The relative abundance of each *Blastocystis* subtype within each concurrent pattern in NHPs.**

**Figure S3. The associations between the gut microbiome and *Blastocystis* load across different patterns in *M. mulatta*. (a)** Absolute abundance of *Blastocystis* across each concurrent pattern. Log2-transformed values are displayed on the Y-axis. **(b)** The Shannon index across subtype concurrent patterns. **(c)** The Principal Coordinates Analysis (PCoA) of the gut microbiota at the species level, based on Bray-Curtis distances. **(d)** PCoA of the gut microbiome enterotypes. **(e)** The relative abundances of Ruminococcaceae and *Limosilactobacillus* within the identified enterotypes. The analysis for panels c and d was adjusted for confounding variables, including sex, age, and BMI. The *P* values were determined using PERMANOVA analysis for panel c, and the Wilcoxon rank-sum test for panels, a, b, and e.

## Supplementary Tables

**Table S1. The *Blastocystis* subtypes and OTU numbers in amplicon sequencing analysis of NHPs.**

**Table S2. The correlation between the absolute abundance of *Blastocystis* and each taxon in *M. fascicularis.*** The analysis was performed using MaAsLin2. Only results for the taxa with a prevalence > 0.1 and q value < 0.25 after *FDR* correction are shown.

## References

1. Piperni, E. et al. Intestinal Blastocystis is linked to healthier diets and more favorable cardiometabolic outcomes in 56,989 individuals from 32 countries. Cell 187, 4554–4570.e18 (2024).

2. Beghini, F. et al. Large-scale comparative metagenomics of Blastocystis, a common member of the human gut microbiome. ISME J. 11, 2848–2863 (2017).

3. El Safadi, D., et al. Children of Senegal River Basin show the highest prevalence of Blastocystis sp. ever observed worldwide. BMC Infect. Dis. 14, 164 (2014).

4. Poirier, P., Wawrzyniak, I., Vivarès, C. P., Delbac, F. & El Alaoui, H. New insights into Blastocystis spp.: a potential link with irritable bowel syndrome. PLoS Pathog. 8, e1002545 (2012).

5. Cifre, S. et al. Blastocystis subtypes and their association with Irritable Bowel Syndrome. Med. Hypotheses 116, 4–9 (2018).

6. Deng, L., Wojciech, L., Gascoigne, N. R. J., Peng, G. & Tan, K. S. W. New insights into the interactions between Blastocystis, the gut microbiota, and host immunity. PLoS Pathog. 17, e1009253 (2021).

7. Tito, R. Y. et al. Population-level analysis of Blastocystis subtype prevalence and variation in the human gut microbiota. Gut 68, 1180–1189 (2019).

8. Andersen, L. O., Bonde, I., Nielsen, H. B. & Stensvold, C. R. A retrospective metagenomics approach to studying Blastocystis. FEMS Microbiol. Ecol. 91, 1–9 (2015).

9. Asnicar, F. et al. Microbiome connections with host metabolism and habitual diet from 1,098 deeply phenotyped individuals. Nat. Med. 27, 321–332 (2021).

10. Stensvold, C. R. & Clark, C. G. Pre-empting Pandora’s box: Blastocystis subtypes revisited. Trends Parasitol. 36, 229–232 (2020).

11. Stensvold, C. R. & Clark, C. G. Current status of Blastocystis: A personal view. Parasitol. Int. 65, 763–771 (2016).

12. Köster, P. C. et al. Intestinal protists in captive non-human primates and their handlers in six European zoological gardens. Molecular evidence of zoonotic transmission. Front. Vet. Sci. 8, 819887 (2021).

13. Scanlan, P. D., Stensvold, C. R. & Cotter, P. D. Development and application of a Blastocystis subtype-specific PCR assay reveals that mixed-subtype infections are common in a healthy human population. Appl. Environ. Microbiol. 81, 4071–4076 (2015).

14. Deng, L. et al. Colonization with two different Blastocystis subtypes in DSS-induced colitis mice is associated with strikingly different microbiome and pathological features. Theranostics 13, 1165–1179 (2023).

15. Deng, L. et al. Colonization with ubiquitous protist Blastocystis ST1 ameliorates DSS-induced colitis and promotes beneficial microbiota and immune outcomes. NPJ Biofilms Microbiomes 9, 22 (2023).

16. Deng, L. & Tan, K. S. W. Interactions between Blastocystis subtype ST4 and gut microbiota in vitro. Parasit Vectors 15, 80 (2022).

17. Yason, J. A., Liang, Y. R., Png, C. W., Zhang, Y. & Tan, K. S. W. Interactions between a pathogenic Blastocystis subtype and gut microbiota: in vitro and in vivo studies. Microbiome 7, 30 (2019).

18. Lepczyńska, M. & Dzika, E. The influence of probiotic bacteria and human gut microorganisms causing opportunistic infections on Blastocystis ST3. Gut Pathog. 11, 6 (2019).

19. Scanlan, P. D. & Stensvold, C. R. Blastocystis: getting to grips with our guileful guest. Trends Parasitol. 29, 523–529 (2013).

20. Maloney, J. G., Molokin, A. & Santin, M. Next generation amplicon sequencing improves detection of Blastocystis mixed subtype infections. Infect. Genet. Evol. 73, 119–125 (2019).

21. Rojas-Velázquez, L. et al. Use of next-generation amplicon sequencing to study Blastocystis genetic diversity in a rural human population from Mexico. Parasit. Vectors 12, 566 (2019).

22. Maloney, J. G., Molokin, A., da Cunha, M. J. R., Cury, M. C. & Santin, M. Blastocystis subtype distribution in domestic and captive wild bird species from Brazil using next generation amplicon sequencing. *Parasite Epidemiol*. Control 9, e00138 (2020).

23. Forsell, J. et al. The relation between Blastocystis and the intestinal microbiota in Swedish travellers. BMC Microbiol. 17, (2017).

24. Tan, K. S. W. New insights on classification, identification, and clinical relevance of Blastocystis spp. Clin. Microbiol. Rev. 21, 639–665 (2008).

25. Karamati, S. A. et al. Comprehensive study of phenotypic and growth rate features of Blastocystis subtypes 1-3 and 6 in symptomatic and asymptomatic subjects. Iran. J. Parasitol. 14, 204–213 (2019).

26. Scicluna, S. M., Tawari, B. & Clark, C. G. DNA barcoding of blastocystis. Protist 157, 77–85 (2006).

27. Poirier, P. et al. Development and evaluation of a real-time PCR assay for detection and quantification of blastocystis parasites in human stool samples: prospective study of patients with hematological malignancies. J Clin Microbiol 49, 975–983 (2011).

28. Koehler, A. V. et al. Use of a bioinformatic-assisted primer design strategy to establish a new nested PCR-based method for Cryptosporidium. Parasit Vectors 10, 509 (2017).

29. Lalle, M. et al. Genetic heterogeneity at the beta-giardin locus among human and animal isolates of Giardiaduodenalis and identification of potentially zoonotic subgenotypes. Int J Parasitol 35, 207–213 (2005).

30. Cacciò, S. M., Beck, R., Lalle, M., Marinculic, A. & Pozio, E. Multilocus genotyping of Giardia duodenalis reveals striking differences between assemblages A and B. Int J Parasitol 38, 1523–1531 (2008).

31. Buckholt, M. A., Lee, J. H. & Tzipori, S. Prevalence of Enterocytozoon bieneusi in swine: an 18-month survey at a slaughterhouse in Massachusetts. Appl Environ Microbiol 68, 2595–2599 (2002).

32. Li, W. et al. Cyclospora papionis, Cryptosporidium hominis, and human-pathogenic Enterocytozoon bieneusi in captive baboons in Kenya. J Clin Microbiol 49, 4326– 4329 (2011).

33. Kleina, P., Bettim-Bandinelli, J., Bonatto, S. L., Benchimol, M. & Bogo, M. R. Molecular phylogeny of Trichomonadidae family inferred from ITS-1, 5.8S rRNA and ITS-2 sequences. Int J Parasitol 34, 963–970 (2004).

34. Cannon, M. V. et al. A high-throughput sequencing assay to comprehensively detect and characterize unicellular eukaryotes and helminths from biological and environmental samples. Microbiome 6, 195 (2018).

35. Bowles, J., Blair, D. & McManus, D. P. Genetic variants within the genus Echinococcus identified by mitochondrial DNA sequencing. Mol Biochem Parasitol 54, 165–173 (1992).

36. Santín, M., Gómez-Muñoz, M. T., Solano-Aguilar, G. & Fayer, R. Development of a new PCR protocol to detect and subtype Blastocystis spp. from humans and animals. Parasitol Res 109, 205–212 (2011).

37. Chen, S., Zhou, Y., Chen, Y. & Gu, J. fastp: an ultra-fast all-in-one FASTQ preprocessor. Bioinformatics 34, i884–i890 (2018).

38. Magoč, T. & Salzberg, S. L. FLASH: fast length adjustment of short reads to improve genome assemblies. Bioinformatics 27, 2957–2963 (2011).

39. Edgar, R. C. UPARSE: highly accurate OTU sequences from microbial amplicon reads. Nat Methods 10, 996–998 (2013).

40. Minh, B. Q., et al. Corrigendum to: IQ-TREE 2: New models and efficient methods for phylogenetic inference in the genomic era. Mol. Biol. Evol. 37, 2461 (2020).

41. Magdalena, L. Fuzzy rule-based systems. in Springer Handbook of Computational Intelligence 203–218 (Springer Berlin Heidelberg, Berlin, Heidelberg, 2015).

42. Blanco-Míguez, A. et al. Extending and improving metagenomic taxonomic profiling with uncharacterized species using MetaPhlAn 4. Nat. Biotechnol. 41, 1633–1644 (2023).

43. Mu, W. et al. Taeniasis impacts human gut microbiome composition and function. ISME J. 18, (2024).

44. Mallick, H. et al. Multivariable association discovery in population-scale meta-omics studies. PLoS Comput. Biol. 17, e1009442 (2021).

45. Shi, Y., Zhang, L., Do, K.-A., Peterson, C. B. & Jenq, R. R. aPCoA: covariate adjusted principal coordinates analysis. Bioinformatics 36, 4099–4101 (2020).

